# Modeling Montbeillard’s height data of a human male

**DOI:** 10.1101/2025.03.02.641023

**Authors:** I S Shruti, S Vijay Prakash

**Affiliations:** Independent Researchers, Alappuzha, Kerala, India

## Abstract

Growth is a dynamic activity of simultaneous biological processes happening at multiple time-scales varying from orders of fractions of a second to several years. Rather than modeling growth with differential equations, this multiple time-scale dynamics is modeled using a simpler algebraic approach that involves continued fraction of the linear time scale. This algebraic approach offers models that are infinitely differentiable like an exponential function but also robust and superposable like linear equations. Thus, unique insights into growth dynamics can be obtained without much need of a calculus background. Growth of bacterial colonies, yeast cultures, *Drosophila* population, mean individual attributes of *Helianthus* and rats have already been modeled using this approach. In this work, we extend the modeling procedure to individual human growth using Montbeillard’s height measurements of his son starting from birth upto almost 18 years of age. Good fits are obtained on the data and growth rates are estimated directly from the model. Thus, this methodology provides generic, flexible, simpler and more interpretable growth models.

## 1 Introduction

Growth is, perhaps, the most primary indicator of life. The dynamics of growth have been studied at multiple time-scales ranging from seconds to even years. The numbers also vary highly from a single living cell to entire populations of millions. It might seem impossible to study such enormous variation with just a single model. Indeed there are several functions available for the study of growth each suitable for a particular branch of study. The logistic growth equation and its modifications are widely used for population growth studies [1]. The Barayani model [2] is available for the study of bacterial growth. The growth of attributes such as mean height of individuals (*Helianthus*) have been studied with functions borrowed from autocatalytic kinetics [3]. Turner’s generalised model is given for mean weight of mammals [4]. Recently, the *a* − *m* model has been used to model growth in all these cases [5, 6, 7]. The growth in height of human populations has been modeled using the Quadratic-Exponential-Pubertal-Stop (QEPS) growth model which is a combination of four different functions [8]. The Phenomenological Universalities (PUN) approach has been used for modeling the growth of an individual male human [9].

The French naturalist Buffon gave what may be considered as the oldest diligently recorded data of an individual human’s growth in his 36 volume book *Histoire Naturelle* published between 1749 and 1789 [10]. The data was contributed by his friend Montbeillard who took semi-annual height measurements of his son Francois for nearly 18 years starting from 1759. The data given in a tabular form by Buffon was later quoted by Belgian mathematician Quatelet in 1835 [12] and 1870 [13] after which it was pretty much forgotten until 1927 when American anatomist Richard Scammon published his work titled ‘The First Seriatum Study of Human Growth’ [11]. In his work, Scammon plotted Montbeillard’s height data as a function of age and generated the first growth curve of an individual human from infancy to 18 years of age. He made several observations from the appearance of the curve such as the accelerated growth from birth till around five years of age and then a slow yet steady growth until the age of 13. After that the growth again accelerates rapidly indicating puberty and then slows down as Francois turns 18 years old. Scammon also compared the individual growth curve to the mean height growth curves belonging to two populations of Parisian children. He observed that both these curves showed similar trends such as the early childhood and pubertal growth spurts [14].

In this work, the *a* − *m* model has been used to model Montbeillard’s data. The advantage of the *a* − *m* model is that it behaves like a line but can be tweaked into a curve with a single parameter *a*.

## 2 Mathematical preliminaries

The linear equation *y* = *mx* with *m* as slope is singularly perturbed as

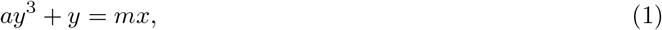

where *a* is a positive parameter. This way the above equation is only parametrically nonlinear but fundamentally linear. We choose *y*^3^ to introduce nonlinearity because the one-to-one correspondence of *x* and *y* of linear equations is preserved. In other words, on an *xy* − plane for every *x* there is a corresponding unique *y* for *a >* 0.

Following Cardano’s method [15] of solving cubic equations, the real solution of Eqn. (1) is a sum of two curves say *S*_1_ and *S*_2_ i.e., *y* = *S*_1_ + *S*_2_, where

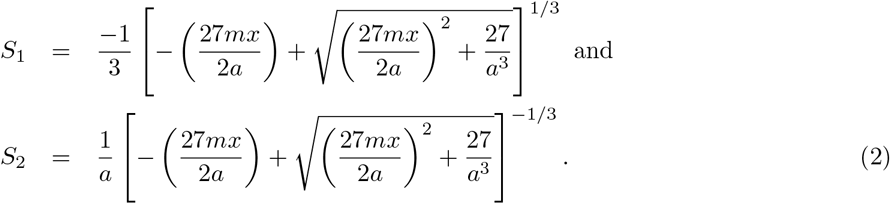

*S*_1_ and *S*_2_ do not intersect on an *xy*− plane [16]. For every *y* there is a unique *S*_1_ and *S*_2_. We can thus trace *y* on an *S*_1_*S*_2_ plane. For *a* → 0, *S*_1_ and *S*_2_ become unbounded and Eqn. (1) becomes *y* = *mx* which is the straight line equation that is conventionally used to span *xy*− plane. Therefore, *xy*− plane is also an unbounded *S*_1_*S*_2_− plane. Eqn. (2) resembles power law expressions [17] with an additional 27*/a*^3^ term within the square root part. This additional term is an adjustment to the power law behavior of *S*_1_ and *S*_2_. Therefore, *a* can be considered as an adjustment parameter. As shown in Fig. (1), for each *a* we get a unique adjusted *xy*− plane. Eqn. (1) can also be expressed as a continued fraction of *y* = *mx* in the following manner

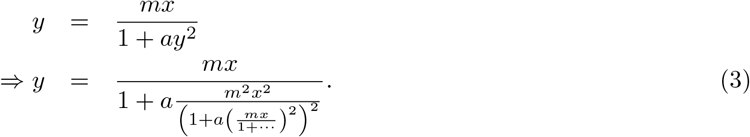

**Figure 1:**
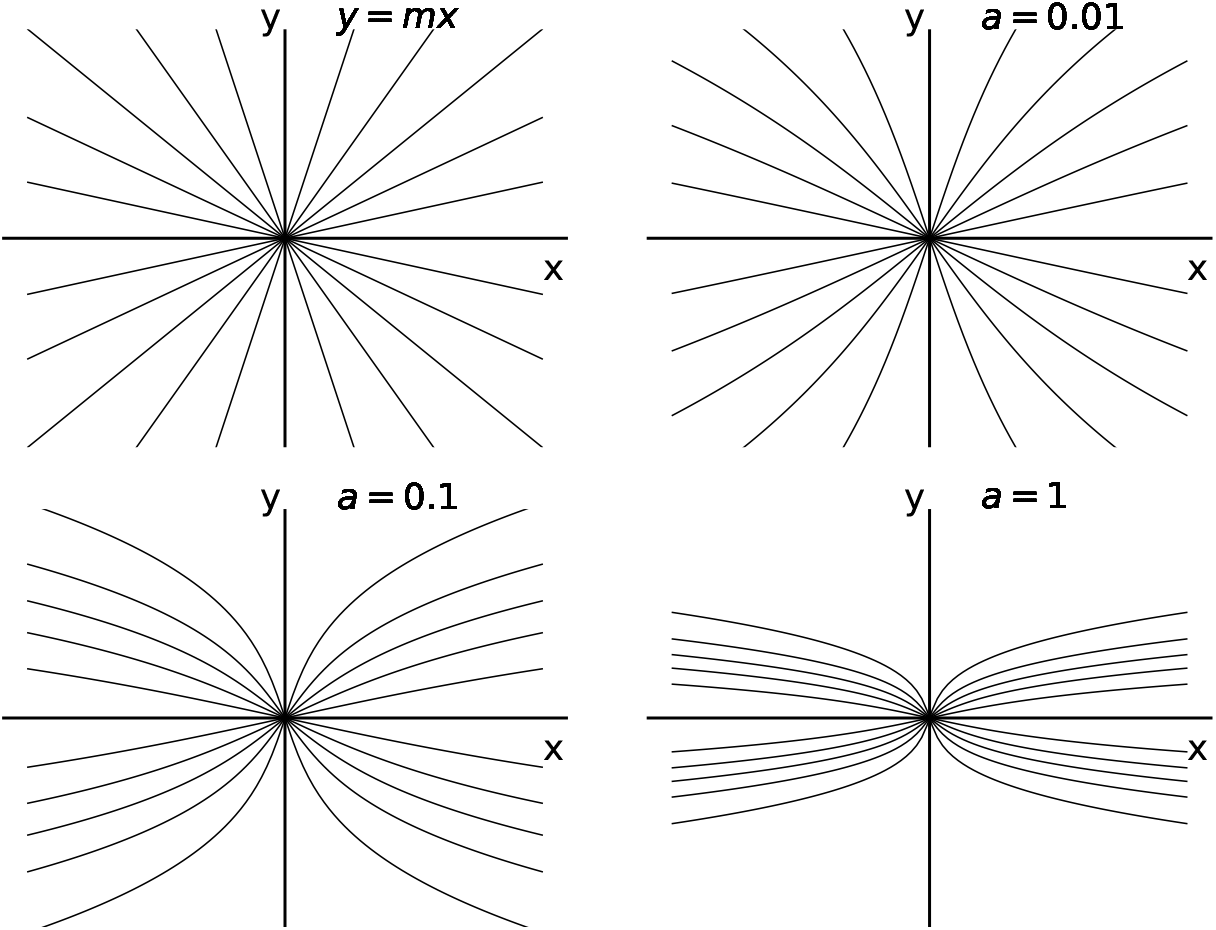
Values of *y* are bounded when *ay*^3^ is introduced to *y* = *mx* equation. Adjusted *xy*− plots for various *a* are shown.

The above equation is only a representation of Eqn. (1) and the latter is the function that is actually used in the fitting procedure. However, this representation showcases how multiple scales of *x* are introduced through *a*. Eqn. (3) obviously converges to the sum *S*_1_ + *S*_2_ as they represent the same *y*. In general, two simultaneous processes characterize multiple scale problems [18]. Here, the simultaneous processes *S*_1_ and *S*_2_ are captured by Eqn. (1) which is also a continued fraction of straight lines.

In the following section we superpose several *a* − *m* models while keeping *a* fixed across models. This is because for any given *a* we can get a uniquely bent i.e. adjusted *xy*− plane as shown in Fig. (1) that models the given set of data points.

### 2.1 Superposition

In this section, we elaborate how the sum of *a* − *m* models can be used to fit nonlinear functions such as the sine and the Gauss error functions. The generalized form of Eqn. (1) is given by

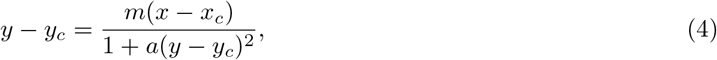

where (*x*_*c*_, *y*_*c*_) is the origin instead of (0,0). The real solution of above equation is given by

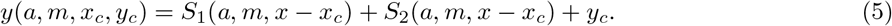

Many of such *y*’s with different origins can be superposed similar to linear equations while fixing *a*. The sum over *n* origins is given by

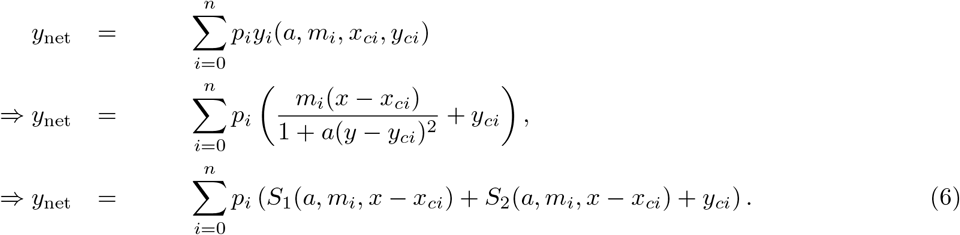

where *p*_*i*_’s are weights, *m*_*i*_ is the *i*^th^ slope with *x*_*ci*_ and *y*_*ci*_ as the origin coordinates. The superposed model can also be written as [7]

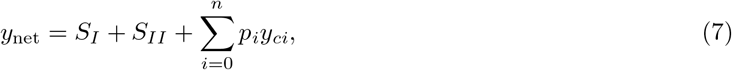

where 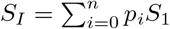 and 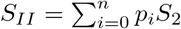.

In the following section, the superposed *a* − *m* models are used to describe sinusoidal waves and the Gauss error function to showcase the usefulness of superposition and the flexibility it offers.

#### 2.1.1 Translation and rotation

Here, following the fitting procedure in [6], we initially fit the sin(*x*) function within the interval [−*π/*2, *π/*2] with superposed *a* − *m* models (as shown in Fig. 2). Thereafter we extend the fit to a wider interval of *x* simply by translation and rotation (as shown in Fig. (3)) similar to the linear equation. This exercise is presented here to showcase how the parametrically nonlinear fit can be used like straight lines. For Fig. (2), the origins are: (0.094, 0.094), (0.031, 0.031), (−0.031, −0.031) and the corresponding slopes *m*_*i*_’s are 3.26, −3.29, 0.5 and *p*_*i*_’s are −1.412, −4.077, −15.7. Thus a weighted sum of lines having slopes with the *m*_*i*_ values given above is used to represent the sine function within the interval [−*π/*2, *π/*2] on an adjusted *xy*− plane with *a* = 1.782 *×* 10^−3^. This sum can be translated and rotated similar to linear equations as shown in Fig.(3).

**Figure 2:**
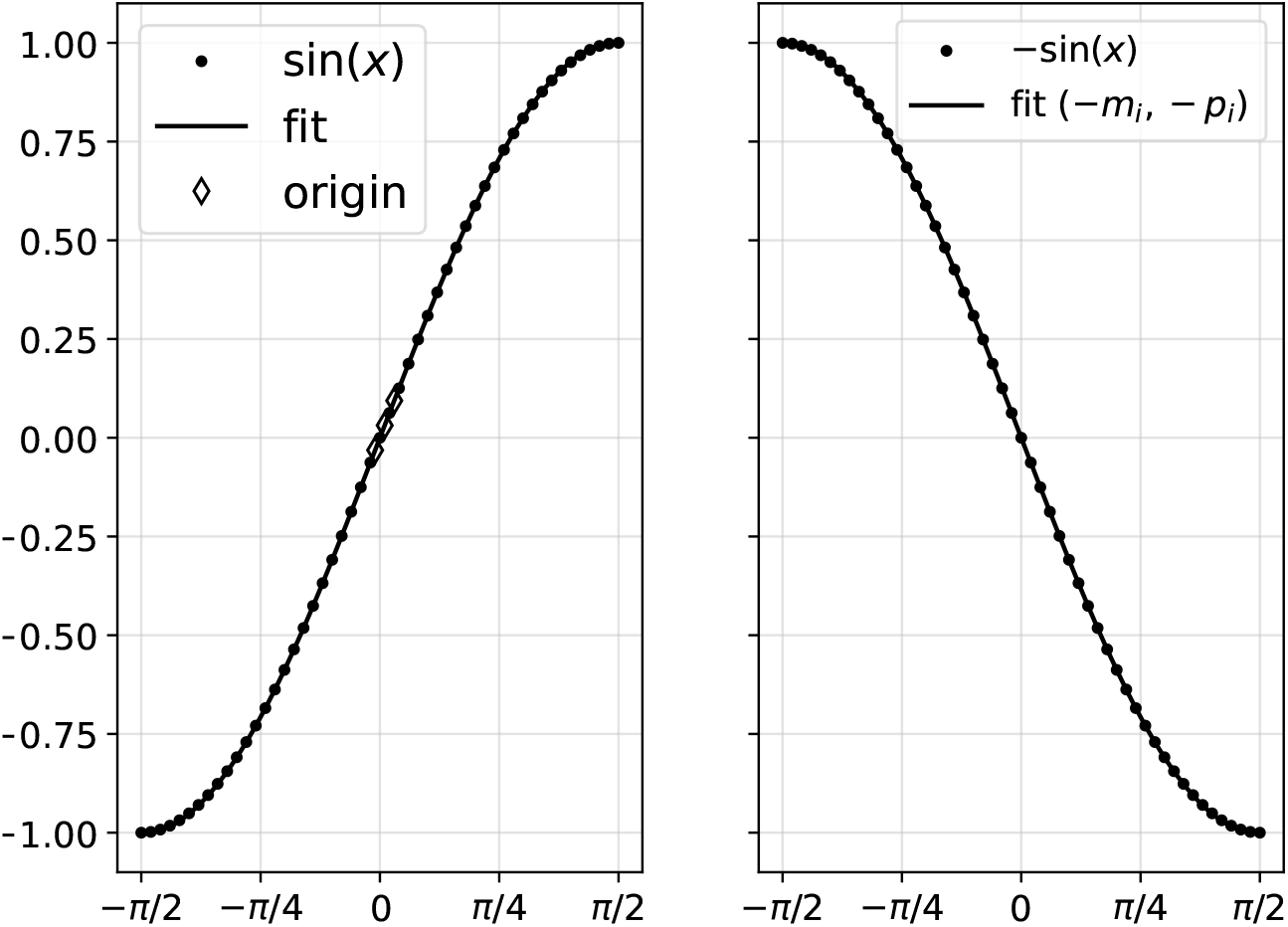
Sine wave within the interval [−*π/*2, *π/*2] on the left hand side. On the right hand side, a simple rotation is carried out by changing the signs of slopes and intercepts similar to linear equation.

**Figure 3:**
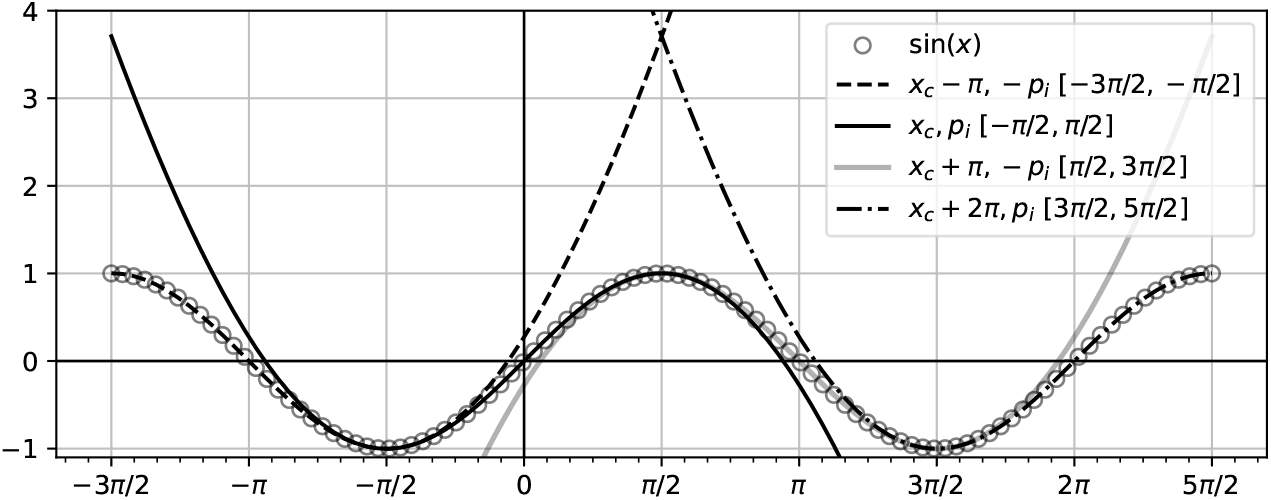
The fit within the interval [−*π/*2, *π/*2] is extended across a wider range of *x* simply by translation through origins and rotation with *p*_*i*_’s.

### 2.1.2 Smoothness

The superposed models are as smooth as exponential functions. This is showcased by fitting the superposed model on Gauss error function. Subsequently, we match their respective derivatives as shown in Fig. (4). Thus, *a* provides a smooth adjustment to the *xy*− plane as shown in Fig. (1). Therefore, while fitting function values their respective derivative information are also taken into account using this model.

**Figure 4:**
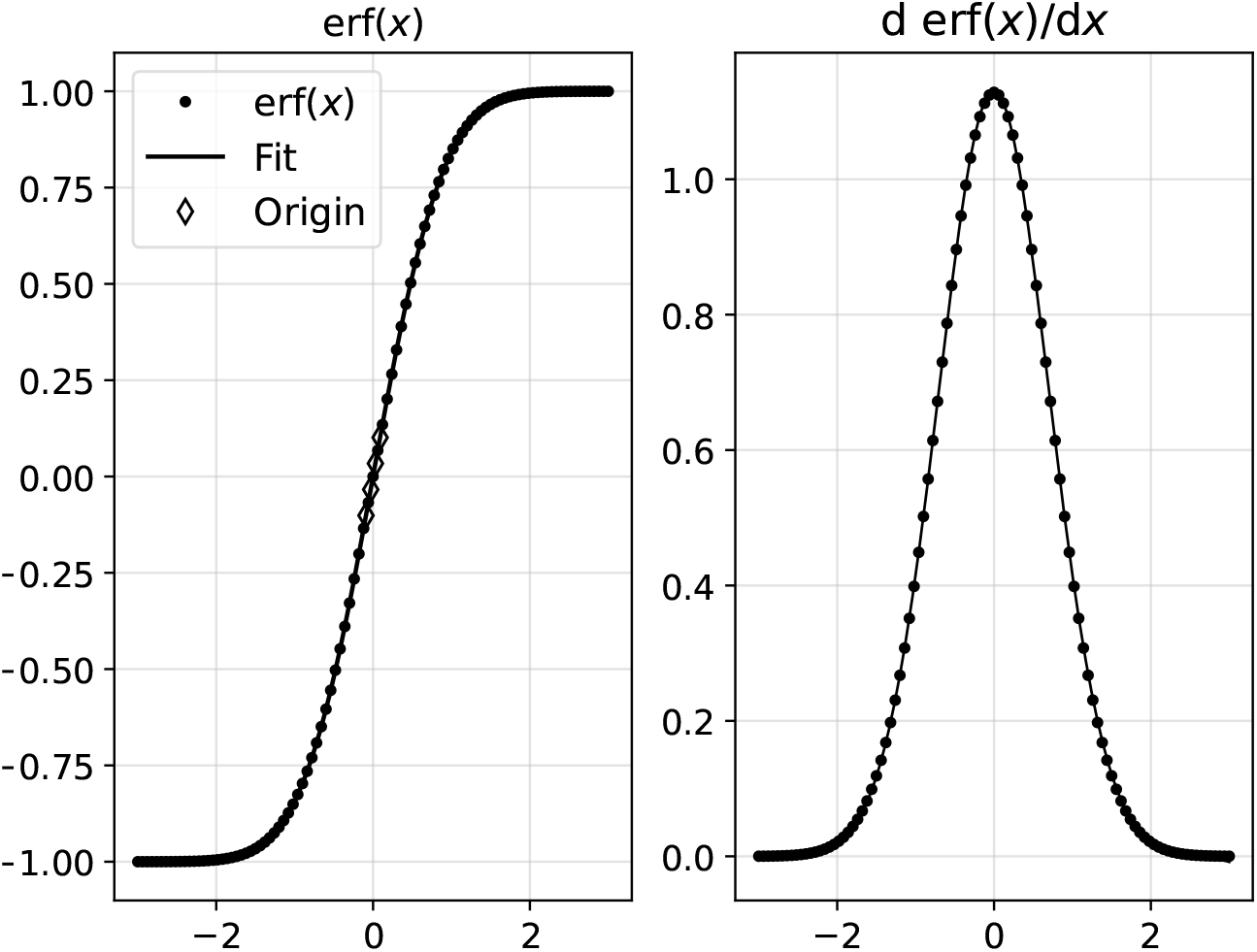
Superposed *a* − *m* models (shown as a continuous line) fit nonlinear error function. Their corresponding derivatives i.e., derivative of the fitted model *dy*_net_*/dx* and the derivative of the error function 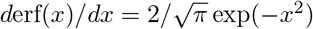 also match.

Here, *a* = 8.0825 *×* 10^−2^, chosen origins are (−0.09, − 0.1), (0.09, 0.1), (−0.03, −0.034), (0.03, 0.033) with corresponding slopes −0.636, 0.636, 0.635, −1.174 *×* 10^−4^ and *p*_*i*_’s given by 3090.04, −1544.43, 4644.47, 18511.01.

### 2.2 Highlights

The following are notable highlights of using *a* − *m* models as a tool for fitting nonlinear data

1. An *a* − *m* model is a sum of two components *S*_1_ and *S*_2_. Each component resembles a power law relationship with an adjustment using *a*.
2. *S*_1_ and *S*_2_ represent simultaneous processes varying differently in *x*. Thus, the sum of these two components reflects multiple scale nature of data at hand.
3. The sum *S*_1_ + *S*_2_ can also be interpreted as a parametric continued fraction of linear scale as expressed by Eqn. (3) which can be reduced into a series that involves multiple orders of *x* [16]. This can also be realized using Fig. (5) which can be reproduced using the following recursive Python program:

~~~
def fnfr(y,x,a,m,num):
 if num==0:
  return y
 return fnfr(m*x/(1+a*y**2),x,a,m,num-1)
~~~ For generating curves *E*_1_, *E*_2_, *E*_3_ and *E*_4_ we run `[fnfr(m*x,x,a,m,i) for i in [1,2,3,4]]`. Therefore, this interpretation explains how this model encompasses multiple scale variations of *y* with respect to *x*.
4. Similar to linear models, the *a* − *m* models can be superposed, translated and rotated.
5. Each superposed model can be written as a sum of two S-curves: *S*_*I*_ and *S*_*II*_ [7]. *S*_*I*_ and *S*_*II*_ are weighted sums of *S*_1_ and *S*_2_ of each *a* − *m* model, respectively.
6. The superposed *a* − *m* models fit nonlinear functions such as the error function while preserving the derivative information as seen on the right hand side of Fig. (4). Thus, they are suitable for modeling growth and obtaining growth rates.

**Figure 5:**
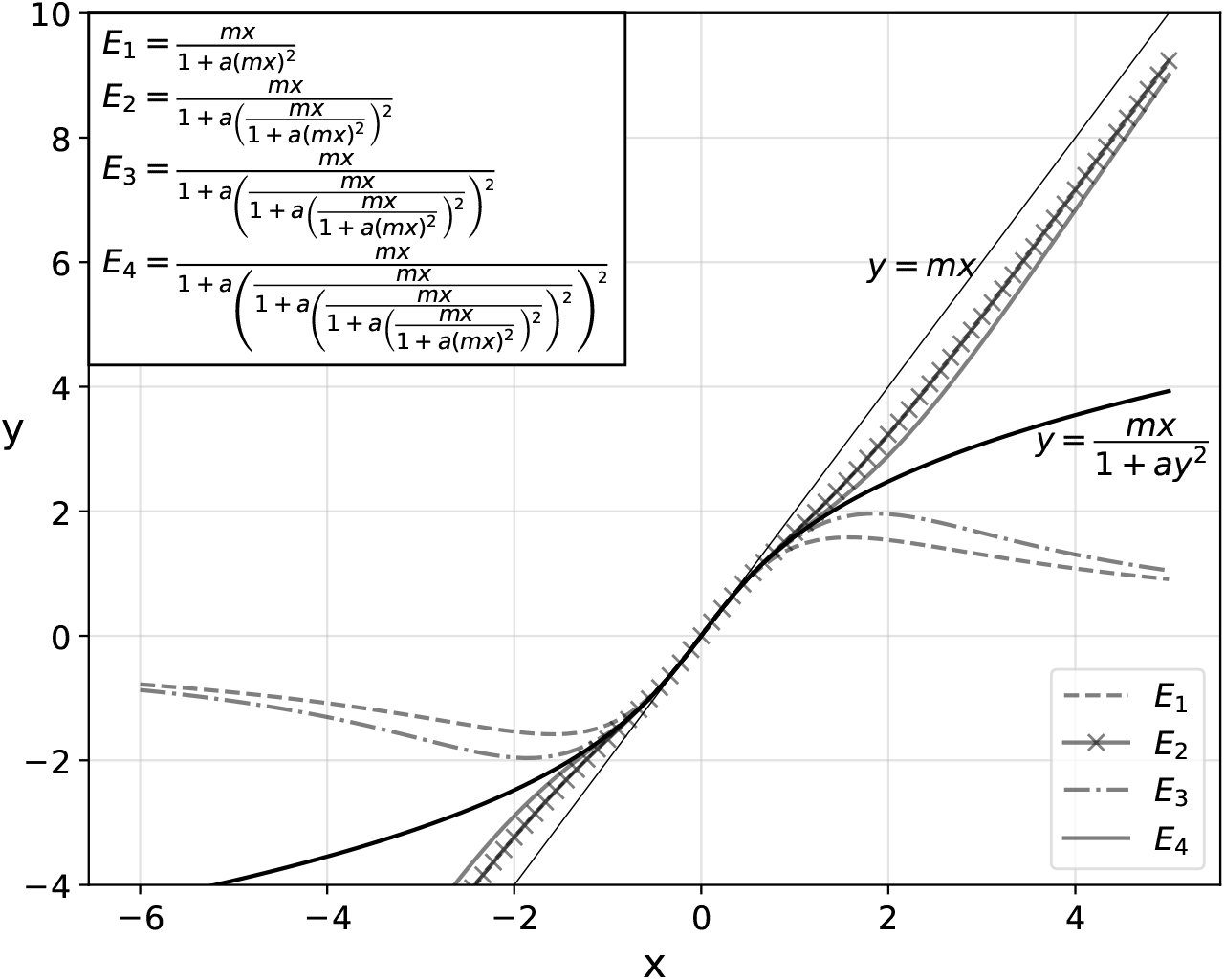
Various levels of continued fraction showing multiple variations with *x* for *a* = 0.1, *m* = 2.

## 3 Growth of a human male

Here, we have applied the *a*− *m* model on Montbeillard’s growth data of his son Francois’ height given in centimeters, measured from when he was born till he is nearly 18. In Fig. (6), as we keep increasing the number of origins the model fits better on the growth data. We then keep increasing the origins and observe the corresponding growth rates. This is shown in Fig. (7). Finally, graphically observable growth rate variations are obtained using the model with 11 origins.

**Figure 6:**
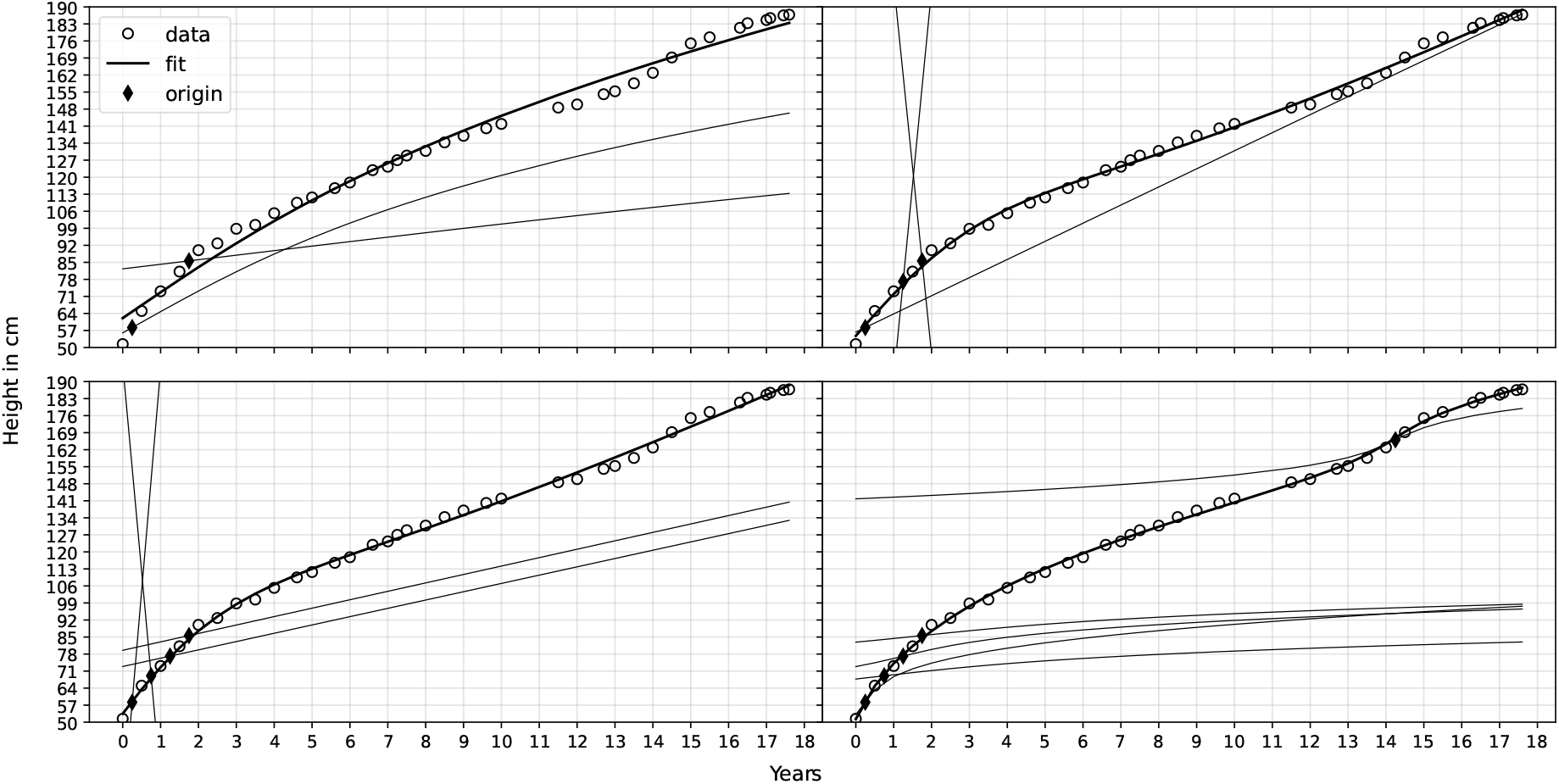
Superposed *a* − *m* models as we increase the number of origins. The plots of individual *a* − *m* models passing through their respective origins are also shown

**Figure 7:**
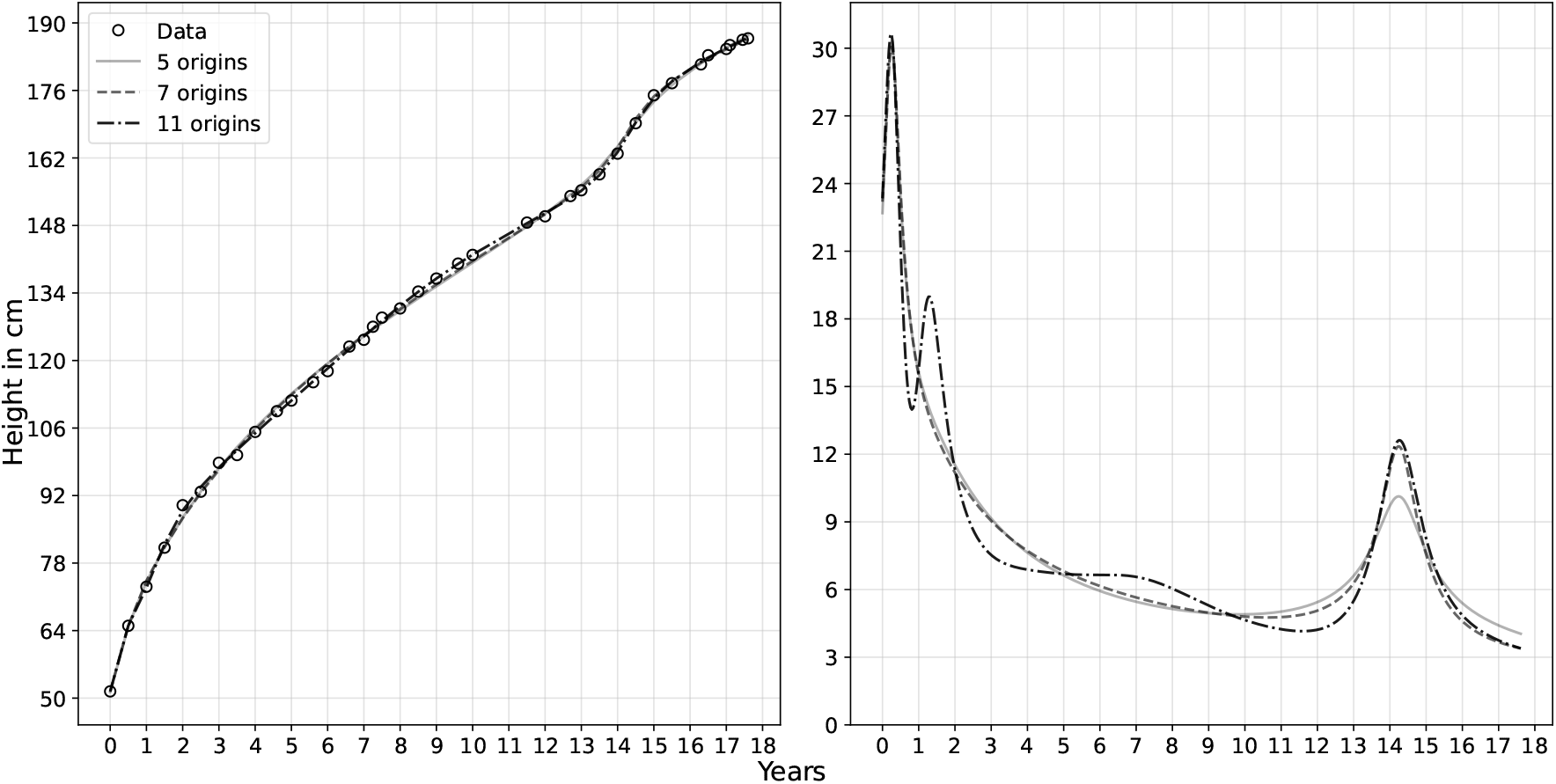
Superposed *a* − *m* models for multiple origins and their corresponding growth rates.

In Fig 8, the curves for *S*_*I*_ and *S*_*II*_ are given as well as the final curve obtained from the 11 origin *a* − *m* model. The final curve is the sum of two simultaneous processes *S*_*I*_ and *S*_*II*_ .

**Figure 8:**
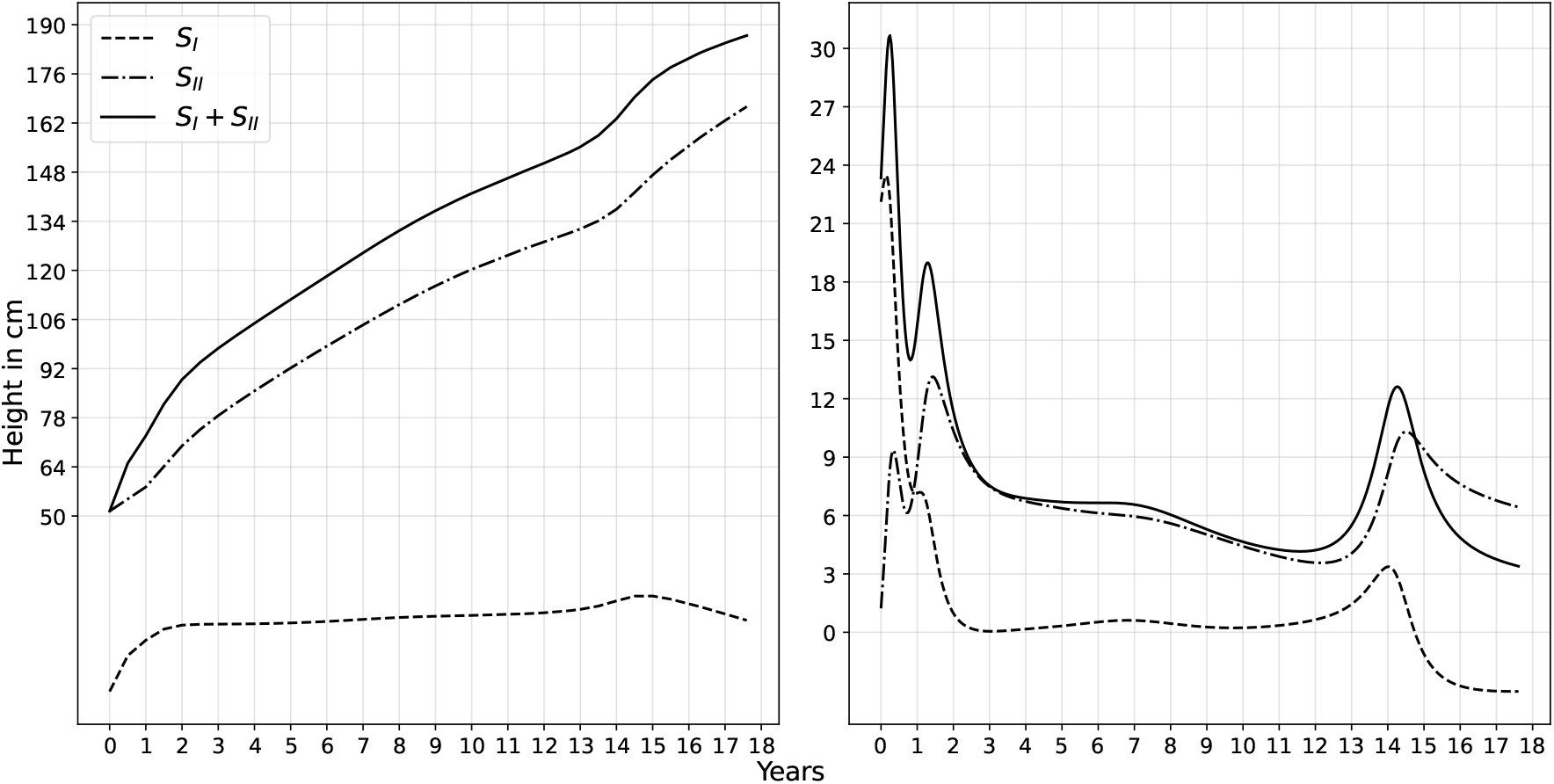
Superposed *a* − *m* models for 11 origins and their components *S*_*I*_ and *S*_*II*_ and their corresponding growth rates.

### 3.1 Observations

1. As can be seen from Fig. (7), the *a*− *m* model with 11 origins fits the data well as compared to the 5 origin and the 7 origin models. This means that the growth is highly nonlinear and each origin represents a significant phase of growth in time.
2. The plots for corresponding growth rates of each model are given on the right hand side in Fig. (7). It can be seen that the growth rate of the 11 origin *a* − *m* model captures two extra peaks of growth around age 2 and 7 years. This means that the 11 origin model is more precise than the others. It can also be interpreted that overall Francois’ growth has occurred in spurts. It is the time taken by these spurts that actually varies. Since the early childhood and pubertal growth spurts happen within a shorter duration, they are more apparent.
3. Using Eqn. (2), the model can be split into two components. Each of these components are shown in Fig. (8).
4. From Fig. (8), it can be seen that *S*_*I*_ shows an increasing trend in the initial years of growth and during puberty while appears to grow almost linearly in between. *S*_*II*_ , on the other hand, shows a slight decline during early childhood and puberty but shows steady increase in between.
5. On the right hand side in Fig. (8), the plots of the corresponding growth rates of *S*_*I*_ and *S*_*II*_ and their sum, the 11 origin *a* − *m* model are given. From the figure it is evident that *S*_*I*_ peaks first followed by *S*_*II*_ while the peak of their sum lies in between. This means that a growth spurt is the sum of successive maxima of two simultaneous processes represented by *S*_*I*_ and *S*_*II*_ .
6. It is also seen from Fig. (8) that when *S*_*I*_ declines *S*_*II*_ increases and vice versa. As Francois approaches 18 years of age, *S*_*I*_ shows a steady decline. *S*_*II*_ too starts declining but only later.

### 3.2 *S*_*I*_*S*_*II*_plane

In this section, we plot the results on an *S*_*I*_*S*_*II*_ coordinate system [7]. Fig. (9) again reinstates that the *S*_*I*_ process shows significant increase during the early childhood and pubertal spurt while the *S*_*II*_ process represents the steady and almost linear growth through out the years.

**Figure 9:**
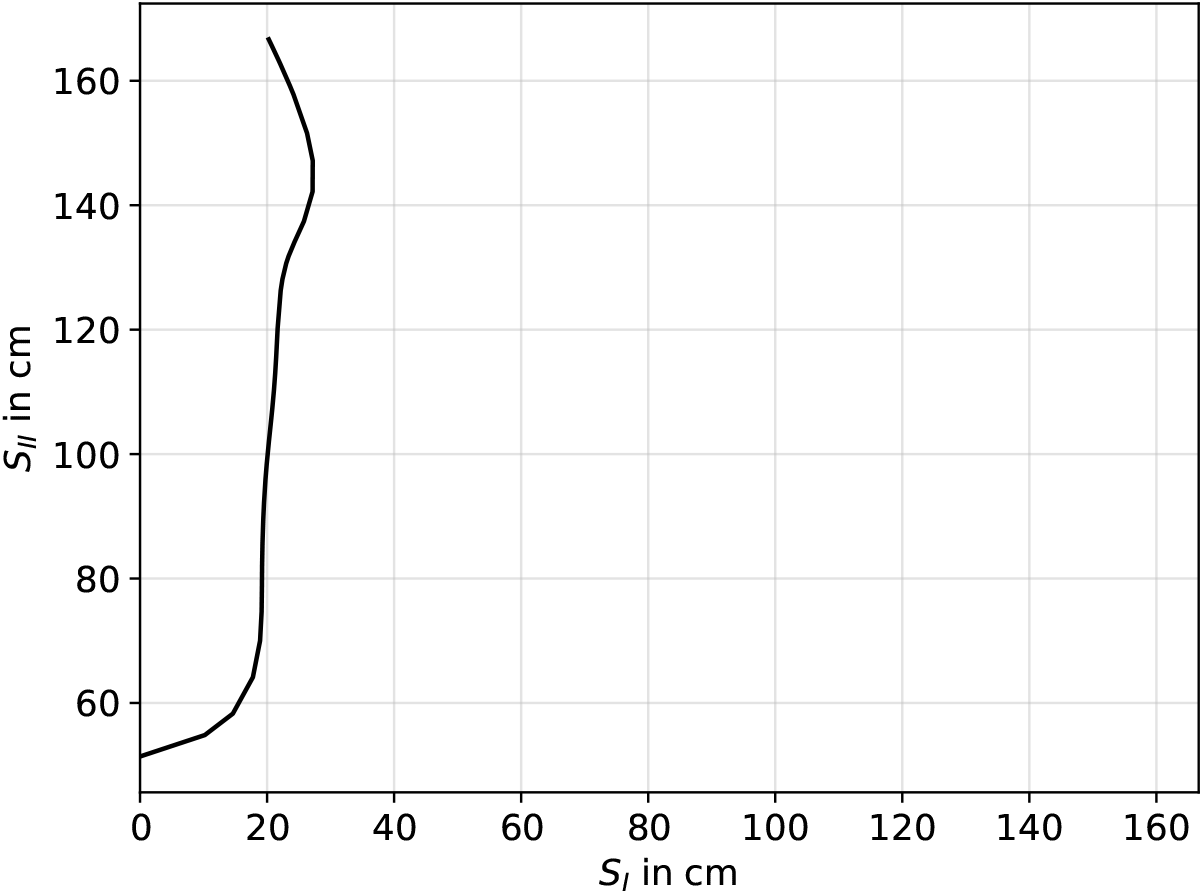
Superposed *a* − *m* models for 11 origins and their components *S*_*I*_ and *S*_*II*_ as axes.

## 4 Conclusions

In this work, we have presented the salient features of the *a* − *m* model and used it to understand the oldest available data representing the growth in height of a male human. The model captures multiple scale nonlinear variations while offering convenient superposed linear forms. As we keep increasing the number of origins the growth rate values converge and we obtain precise estimates of growth rates. Finally we obtained a good fit with 11 origins and derived the corresponding growth rates directly from the fit. It is observed that the overall growth of the individual has occurred in spurts. The duration taken for these spurts varies. The early childhood and pubertal spurts are the most rapid while the mid-childhood growth spurt happens over several years and peaks around the age seven. It is also inferred from the model that the overall growth process is a sum of two simultaneous processes. The interplay of these two processes which may be attributed to intrinsic and extrinsic factors decides the rate of growth and the timing of the growth spurts.

## Notes

### Competing Interest Statement

The authors have declared no competing interest.

